# An artifact-robust framework for measuring tCS effects during stimulation

**DOI:** 10.1101/2025.10.31.685770

**Authors:** Nima Noury, Fabio Damiani, Markus Siegel

**Affiliations:** Department of Neural Dynamics and Magnetoencephalography, Hertie Institute for Clinical Brain Research, University of Tübingen, Germany; Centre for Integrative Neuroscience, University of Tübingen, Germany; MEG Center, University of Tübingen, Germany; Center for Bionic Intelligence Tübingen Stuttgart (BITS), Tübingen, Germany; German Center for Mental Health (DZPG), Tübingen, Germany

**Keywords:** tACS, MEG, artifacts, visual flicker, interaction, brain-state dependency

## Abstract

**Objective:** Transcranial current stimulation (tCS) is a promising technique to non-invasively modulate human brain activity. However, stimulation artifacts in EEG and MEG recordings severely hinder the study of its online effects.

**Approach:** Here, we introduce a new approach to account for these artifacts. The approach rests on two key ideas. First, we focus on interactions of tCS with intrinsic brain activity, which are absent for tCS artifacts. Second, rather than removing artifacts, we compare composite signals that share similar artifacts but potentially differ in the interaction of tCS with intrinsic brain activity. We follow a simple logic: if tCS does not interact with intrinsic brain activity, then neural activity during simultaneous sensory stimulation and tCS should equal the linear superposition of the neural effects of each applied alone. Any deviation from this prediction provides evidence for a neural interaction.

**Main results:** We tested this approach in a proof-of-principle MEG study, applying 10 Hz transcranial alternating current stimulation (tACS) during rest and during a 10 Hz visual flicker. We compared neural activity during simultaneous stimulation with that predicted by the linear superposition of flicker and tACS alone and found a phase-dependent interaction between tACS and flicker-evoked brain activity.

**Significance:** Our work establishes a novel approach to investigate online effects of tCS and suggests a state-dependent interaction of tACS with human brain activity.

## Introduction

Transcranial current stimulation (tCS) is a non-invasive brain stimulation technique, which allows control over stimulation strength, frequency and – to some extent – stimulation site [1–4]. These features render tCS an attractive technique for manipulating specific brain rhythms [5]. The potential of electrical stimulation to modulate neuronal oscillations has been demonstrated in animal models [6–9]. However, despite progress [10,11], stimulation artifacts remain a major challenge for investigating modulations of human brain activity during stimulation, and it has been argued that prior findings may have been contaminated by residual artifacts [12–15].

Here, we propose a novel approach to study online effects of tCS while accounting for stimulation artifacts. Our approach rests on two key ideas. First, behavioral and offline effects of tCS depend on the state of ongoing brain activity, indicating that online effects may arise through interactions of tCS and intrinsic brain activity [6,16–25]. Such interactions are, by definition, absent for mere tCS artifacts. Thus, our approach aims to identify interactions of tCS with intrinsic brain activity. We therefore test for interactions by controlling intrinsic activity with sensory stimulation and asking whether neural activity during simultaneous sensory stimulation and tCS matches the linear superposition of the effects of each applied alone. Any deviation from linear superposition indicates an interaction between tCS and intrinsic activity. Second, because tCS artifacts are non-linear and several orders of magnitude larger than intrinsic signals [12–15] we avoid attempting to subtract artifacts from recorded signals. Instead, we construct and compare composite signals that are matched in artifact level but potentially differ in whether they contain interactions between tCS and intrinsic brain activity.

We implemented this approach in a proof-of-principle magnetoencephalography (MEG) study. We induced a well-controlled 10 Hz brain rhythm using visual flicker (steady-state response) and simultaneously applied 10 Hz transcranial alternating current stimulation (tACS) at different phase relations to the flicker. This allowed us to test whether our approach reveals neural interactions between tACS and intrinsic brain activity during stimulation. We observed clear deviations from the linear prediction, indicating phase-dependent interactions between tACS and intrinsic brain activity.

Manuscript structure: The Results begin with a concise, intuitive overview of the experimental approach, followed by the proof-of-principle MEG experiment and its outcomes. Complete methodological details are provided in Materials and Methods and can be consulted as needed.

## Materials and Methods

### Participants and experimental protocol

Sixteen healthy subjects participated in this study. All subjects provided written consent and were financially compensated. The study was carried out in accordance with the Declaration of Helsinki and was approved by the local ethics committee. None of the subjects reported a history of psychiatric/neurological disorders or any other contraindications of tACS.

Each subject participated in six blocks of 280 trials. Seven distinct conditions were applied (Fig. 2). These conditions were arranged in randomized seven-trial sequences such that each condition was applied once before a new seven-trial sequence began. This resulted in 40 trials per condition per block. Of the seven conditions, one condition only contained a 10 Hz visual flicker stimulus (V), and two contained only 10 Hz tACS, one with an initial phase of 0 (T_0_) and the other one with an initial phase of *π* (T_π_). The remaining 4 conditions contained simultaneous 10 Hz visual flicker and 10 Hz tACS, and each had a phase difference of either 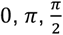, or 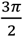 between the two (VT_0_, VT_π_, VT_π /2_, 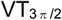, respectively). VT_0_ and VT_π_ were constructed by simultaneous application of V and T_0_ or T_π_ conditions, respectively. To construct VT_π /2_ and 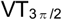 conditions, we also combined V and T_0_ or T_π_ conditions but started V signals 25 ms (i.e. 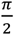 radians at 10 Hz) earlier than for the original V condition (Fig. 2). We used signals after tACS offset of T_0_ and T_π_ conditions as rest signals (R, details explained below).

Each trial was 1.2 sec long, and was followed by an inter-trial interval (ITI) with a random length that was uniformly distributed from 1 to 1.4 sec. During all conditions, subjects were required to simply focus on a white fixation point. After every second inter-trial interval, the fixation dot changed from white to blue for one second. Subjects were asked to restrict their eye blinking to this interval. This helped us to minimize blinking artifacts.

### Visual flicker stimulus

Visual stimuli were presented on a back-projection screen using a projector with 60 Hz refresh rate. The visual flicker stimulus was constructed of 10 flashes. Each flash was 1 frame long and successive flashes were 6 frames apart from each other, which yields a visual flicker at 10 Hz. On each flash, a white narrow horizontal rectangle on a black background at full contrast was shown. We chose this shape to optimize the simultaneity of the retinal input because the projector rendered each frame in horizontal lines from top to bottom. Specifically, we set the height of the rectangle such that there was a 2 ms delay between the onset of the top and bottom edges of the white rectangle (i.e., 7.2 degrees at 10 Hz). A fixation dot was shown during the entire experiment. During flashes, we presented the fixation dot on a small black circle. The fixation dot was white, except for eye-blinking intervals, during which it turned blue. Stimulus presentation was designed and controlled via custom scripts using Psychtoolbox-3 [26]. After each experiment, we checked the timing of the presented frames as reported by Psychtoolbox and did not observe any frame drops or refresh rate distortions. Moreover, we used a photodiode to further verify the timing of the visual flicker stimulus (explained below) and did not observe any distortion in the 10 Hz visual flicker.

### Transcranial current stimulation

The stimulation current was applied with an IZ2H stimulator (Tucker Davis Technologies Inc.) driven by an LZ48-400 battery pack. The 10 Hz tACS signal was generated by an RZ5D BioAmp Processor and was controlled via processing chains that were created using the RPvdsEx Visual Design software. These processing chains were activated using custom-written Matlab scripts, which made use of the ActiveX software framework to send commands to the signal generator. Stimulation was applied via two 35 cm^2^ rubber electrodes (neuroConn GmbH), which were attached using Ten20 conductive paste (Weaver and Company).

The electrodes were positioned such that the top edge of the first electrode lay over the midline parietal location (PZ) and the top edge of the second electrode lay over the inion (IZ). Cables ran over the right shoulder. This montage was chosen to ensure that the spatial focus of the induced electric field would be close to primary visual cortex. This was validated using SimNIBS [27] (Fig. 2).

In each trial, the stimulation current was linearly ramped up from zero to the maximum intensity over a period of 250 ms to minimize the possibility of unpleasant tactile sensations. At the end of each trial, stimulation stopped at zero amplitude after precisely 12 cycles.

The maximum value of the tACS current was determined for each subject as follows: Electrodes were attached to the chosen scalp locations using conductive gel. Starting at 0.1 mA (peak amplitude), tACS was applied for approximately ten cycles. The intensity was then increased by 0.05 mA without the subject’s knowledge. This incremental increase continued until the subject reported either phosphene perception or any tactile sensation at the electrode locations. At this point the current was decreased by 0.05 mA and the magnitude was recorded for future reference. The electrodes were then secured with gauze and tape, after which the procedure was repeated in a darkened room, to ensure that any phosphene perception would be more pronounced. This time, tACS application began at the previously recorded threshold, and was incrementally increased without the subjects’ knowledge after ten cycles, as before. As soon as phosphene perception was reported, the intensity was decreased and the current was applied again for ten cycles. Depending on feedback from the subject, the intensity would either be increased or decreased. This gradual adjustment continued until the subject reported that no phosphenes were visible. The subject-specific tACS intensity used during the experiment was ultimately chosen to be 0.05 mA lower than the maximum current at which no phosphenes were perceived. The procedure resulted in subject-specific intensities ranging from 0.25 mA to 0.9 mA (stimulation intensity of all subjects in mA: .30, .40, .40, .45, .90, .30, .30, .50, .30, .45, .30, .30, .60, .30, .40, .25. Note that peak-to-peak values would be two times bigger).

We corrected the difference in the internal clock of the tACS stimulator and the clock of the visual flicker computer as follows. We used a photodiode on the monitor to record the exact timing of the visual flicker and recorded its output with the A/D converter of the MEG recording console. Moreover, using the same A/D converter we recorded the tACS stimulation current by measuring the voltage drop across a 200 Ω resistor positioned in series to the head. Next, we carefully adjusted the frequency of the tACS signal until the frequency of the 2 measured signals matched. In addition, due to the response time of the projector, we found a delay of approximately 30 ms between the execution of the Matlab command for flicker presentation and the appearance of the flicker on the projector screen. This was corrected by introducing an extra time delay before tACS onset at the beginning of each trial, which again was determined by careful inspection of the photodiode and tACS voltage traces.

### Data acquisition and preprocessing

We recorded 272-channel MEG (Omega 2000, CTF Systems) throughout the experiment at 2343.8Hz sampling rate. All signals were in the dynamic range of the recording system and no clipping was observed. Due to the interference between stimulation currents and electrical currents of the head-positioning circuits of the MEG system, we could not monitor head movement continuously during the experiment. We recorded ECG signals using bipolar channels of the MEG system. The ECG was recorded through 2 electrodes placed below the right clavicle and below the left pectoral muscle.

We manually inspected all data and removed all parts of the data with physiological or technical artifact. We defined the start of trials as the start of the flicker stimulus in the V condition, and as the start of the tACS stimulus in all other conditions. Then, we segmented the data into 2.4 s intervals, starting 200 ms before trial onset, and linearly detrended each interval (removing a linear fit from each interval, Matlab detrend-function).

We generated time-shifted copies of trials from the V condition, which in the following we term 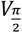. This was done by shifting these trials 25 ms (i.e. 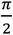 radian in 10Hz) backward in time. These copies were later used for constructing the composite signals.

Because of the difference between the clocks of the MEG recording console and of the tACS stimulator, the tACS frequency appears slightly different from 10 Hz in MEG recordings. To correct this, we manually selected several intervals of MEG data with stable tACS artifacts from the first recorded subject, and estimated their stimulation frequency by fitting a sine wave to each interval (Matlab nlinfit-function), and took the median of the estimated stimulation frequencies (10.000711273294087 Hz). Next, we redefined the time of all MEG data (Matlab interp1-function) to place the recorded stimulation frequency at exactly 10 Hz. Moreover, we resampled the data to 2360 Hz to ensure that the sampling rate was divisible by the stimulation frequency. This helped, at later stages, to minimize the spectral leakage and to obtain FFT estimates exactly at 10 Hz, i.e. at the frequency of interest.

To prevent temporal leakage of stimulation artifacts to the post-stimulation interval, we did not apply any filtering to the data. However, to improve estimates of spectral content at 10 Hz, it was necessary to remove slow drifts of data by detrending. Detrending decreases the spectral leakage from lower frequencies to 10 Hz. To this end, we first cut two 850 ms long segments (i.e. 8.5 cycles at 10Hz) out of each 2.4 s interval, one segment from 350 ms after start of the trial to the end of stimulation, and the other segment starting 150 ms after the end of the stimulation. We refer to the former as during-stimulation segments (*y*_*durin*g_), and to latter as post-stimulation segments (*y*_*post*_). We chose this exact length to improve the performance of linear de-trending for segments with strong tACS artifacts. This is because, the artifact phase at MEG channels with highest tACS artifacts is either close to 0 or 180 degrees [14]. Because a sine wave with a phase near 0° or 180° is temporally symmetric over a duration of 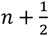 cycles, linear detrending does not distort it or introduce artifacts. Next, we linearly detrended each during-stimulation and post-stimulation segment and discarded the first 50 ms of each segment.

In sum, preprocessing resulted in two segments of data for each trial. Both segments were exactly 800 ms long (i.e. 8 cycles at 10Hz). The during-stimulation segments extended form 400 ms after start of the stimulation to the end of the stimulation. The post-trial segments started 200 ms after stimulation offset. Moreover, the data were resampled in a way that the tACS frequency was recorded at exactly 10 Hz, and that applying FFT to them would provide a spectral estimate at exactly 10 Hz.

For each block (280 trials, i.e. 40 trials per condition), we concatenated all during- and post-stimulation pieces, calculated the beamforming spatial filter, and projected the data to source-space. We used adaptive linear spatial filtering (beamforming) [28] with zero regularization factor (λ) [12]. To get one estimate at each source location, we calculated three orthogonal filters (one for each spatial dimension), projected the data through them, and linearly combined them in the direction of the maximum variance.

### Source locations and physical forward model

We performed the beamforming analysis on a regular three-dimensional grid that covered an SPM’s template brain with 1-cm spacing in MNI space (4805 source locations, spm8/templates/T1.nii). We used fieldtrip’s template T1-weighted structural MRI for all subjects (template/headmodel/standard_mri.mat) and nonlinearly transformed source locations into head space. The MEG sensors were aligned to the head geometry on the basis of three fiducial points (nasion, left ear, right ear) that were registered before and after the MEG acquisition by three head localization coils. To derive the physical relation between sources and sensors (leadfield), we employed a single-shell head model [29].

### Spectral analysis

We applied FFT to each 800 ms data segment at the source level and took the spectral content at 10 Hz for further analysis.

For each subject and at each voxel, we computed the pairwise phase consistency (PPC) as an unbiased measure of phase locking across trials [30]:

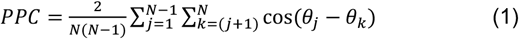

Where *θ* is the phase of each 800 ms segment at 10 Hz and *N* denotes the number of trials. As a measure of power, we calculated the decimal logarithm of the absolute value of FFT results, and took the average across trials, for each voxel and subject.

### Contrasting conditions

We employed cluster permutation statistics to contrast source-level parameters between various conditions (Fig. 3). For each subject, voxel, and condition, we first obtained the mean 10 Hz power across trials (or PPC). Then, at each voxel, we concatenated values of the two compared conditions of all subjects, which resulted in a vector with 32 samples. We normalized this vector by histogram matching (Gaussian), and obtained a zero-mean normal vector, but kept its standard deviation. Next, at each voxel, we took the difference between the two conditions and calculated its t-statistic across subjects. Finally, we employed a cluster sign-flip permutation test on these values. Specifically, we defined spatial clusters of voxels whose t-values exceeded a threshold of 2.581 (or were below −2.581 when testing for negative clusters), using a minimal-neighborhood criterion to determine spatial contiguity. Then, we averaged t-values of all voxels of each cluster. Next, we estimated the null-distribution by randomly flipping the sign of the normalized power (or PPC) differences of each subject (same flip for all voxels of each subject), defining clusters, and keeping the maximum of mean t-values across clusters. We repeated the sign flipping 1000 times and assigned a p-value to each cluster of the original data by comparing its averaged t-value against the distribution of the 1000 maximum values in the permutation distribution.

From the contrast between visual flicker only (V) and rest (R) conditions (Fig. 3d), we defined a region of interest (ROI), in which we subsequently tested for the interaction of tACS and visual flicker, i.e. for cluster permutation tests used to compare composite signals. This ROI was created by combining the power and phase clusters that showed significant differences between the V and R conditions.

### TACS offset artifact

To avoid contamination of post-stimulation segments by residual tACS offset artifacts, we defined these segments to begin 200 ms after tACS offset. Moreover, we did not apply filtering to our data, because any filtering would spread artifacts across time. We performed several tests to ensure that post-stimulation segments did not contain any tACS offset artifacts. As the first test, we argued that tACS conditions that have anti-phase tACS signals, should also show anti-phase tACS offset artifacts. Thus, if post-stimulation segments were contaminated by offset-artifacts, their *PLV*_*complex*_ vectors should point to opposite phases. We tested this using a cluster permutation test, in which we took the difference of *PLV*_*complex*_ vectors of two corresponding tACS conditions and compared the length of the resulting vector against the null-distribution. We constructed the null-distribution by randomly flipping the sign of *PLV*_*complex*_ of subjects (similar to *Contrasting conditions*, but with a threshold of 1.96 to increase the sensitivity). We did not observe any significant difference (T_0_ against T_π_, p>0.65). We performed additional cluster permutation tests to investigate if there was any significant difference between power or PPC of post-stimulation segments of tACS conditions with anti-phase tACS signals. None of these comparisons showed a significant difference (T_0_ against T_π_, P_power_ > 0.3, P_PPC_ > 0.16). In sum, we concluded that the post-stimulation segments were not contaminated by tACS artifacts.

### Composite signals

We added up signals of different conditions to construct composite signals *x* and 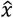 (Fig. 4 and Fig. 5). These composite signals implement the linear superposition logic: if simultaneous stimulation equals the linear prediction, no interaction is present; any deviation indicates a neural effect. We constructed 800 ms long *x* signals by summing during-stimulation segments of the VT conditions with rest segments (R), which were defined as post-stimulation segments of the T conditions (Fig. 2). As there were only two rest conditions but four *x* signals, each rest condition had to be used twice. Therefore, we randomly attributed each rest to one of the *x* signals, as following:

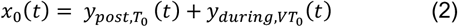

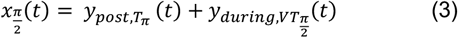

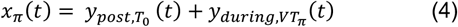

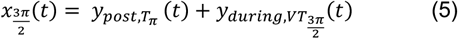

We constructed 800ms 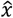 signals by summing up during-stimulation segments of V and T conditions:

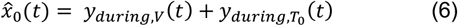

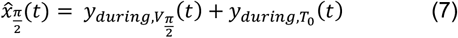

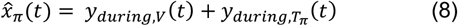

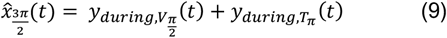

As we were interested in the 10 Hz spectral content, instead of first constructing the *x* and 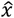 signals in the time domain and then estimating their 10 Hz spectral component, we equivalently first estimated the 10 Hz components of all data segments at the source level and then added up the frequency domain components to estimate the 10 Hz components of *x* and 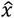.

Furthermore, to quantify power and PPC modulations of *x*_*θ*_ and 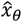 by *θ* (Fig. 5), we constructed corresponding Δ_*θ, θ*+*π*_ and 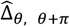 samples, as follows:

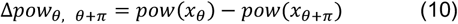

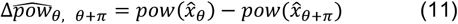

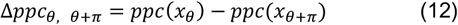

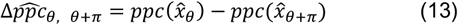

for *θ* = 0 *and* 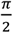.

### Null model and analysis pipeline

First, we compared power and PPC of composite signals against each other (e.g. *x*_0_ against 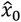, Fig. 4). To this end, we employed the same cluster permutation tests as for power and PPC of regular conditions (see *Contrasting conditions*, we used a threshold of 1.96 for tests on composite signals to increase the sensitivity).

In another cluster permutation test (Fig. 5), we compared |Δ_*θ, θ*+*π*_| with its corresponding 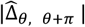, for *θ* = 0 *and* 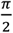, and for power and PPC (|. | refers to absolute value). Given how the composite signals were constructed (equations 2-9), it is clear that *x*_*θ*_ and 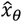 signals at different values of *θ* were not statistically independent. For example, all 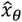 signals used the same trials of *y*_*durin*g,*V*_ (or its time shifted versions). These interdependencies complicate the comparison of |Δ_*θ, θ*+*π*_| and 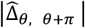 by decreasing the degrees of freedom, which in turn affects the |Δ_*θ, θ*+*π*_| and 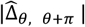 estimates. Importantly, different levels of interdependencies may lead to a spurious difference between |Δ_*θ, θ*+*π*_| and 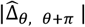. To prevent any false positives due to this issue, we grouped trials into odd and even trials, and constructed new *x*_*θ*_ and 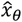 signals based either only on odd trials (*x*_*θ,odd*_ and 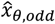) or only on even trials (*x*_*θ,even*_ and 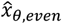).

Next, we used independent pairs of these new composite signals to generate new delta signals:

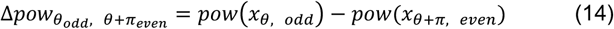

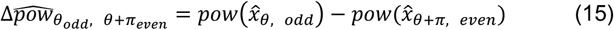

for *θ* = 0 *and* 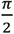 (similarly for PPC). For simplicity, we refer to these as Δ_*θ, odd*_ and 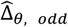. The other option for using independent pairs would be:

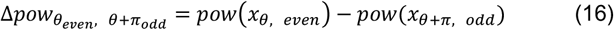

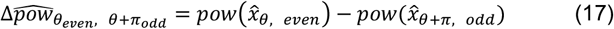

for *θ* = 0 *and* 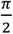 (similarly for PPC). For simplicity, we refer to these as Δ_*θ, even*_ and 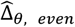. Pay attention that, in each of these new variables, no trial is used twice. Therefore, it is valid to compare |Δ_*θ, odd*_| against 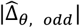, and |Δ_*θ, even*_| against 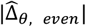. This procedure ensured that any observed differences between composite signals could not arise from statistical dependencies between the constituent trials.

Ideally, one would want to either combine these two tests into one test, or combine their results. Next, we present a method to combine these two tests into one test. We first explain that the presented method leads to asymmetric null-distributions, which complicates the use of permutation tests. Then, we describe a transformation to solve this issue.

One idea to combine the two tests might be to take the average of |Δ_*θ, odd*_| and |Δ_*θ, even*_| (i.e.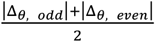), and compare it against the average of 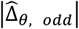 and 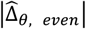 (i.e. 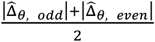, using a sign-flip permutation test. However, such a test would be biased towards false positives, because some of the trials used in |Δ_*θ, odd*_| and 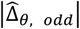, are also used in |Δ_*θ, even*_| and 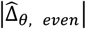 (e.g. looking at equations 6-9, 15 and 17, it is clear that both even and odd trials of *y*_*durin*g,*V*_(*t*) are used in both 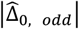 and 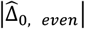). Therefore, |Δ_*θ, odd*_| and |Δ_*θ, even*_|, and also 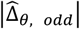 and 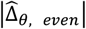 are not independent from each other. Importantly, the amount of statistical dependency in the former pair could be different from the latter pair. Therefore, under the null hypothesis, at each voxel and across subjects, 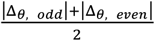 and 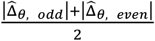 have positively skewed distributions with equal mean but potentially different variances. Hence, under the null hypothesis, at each voxel and across subjects,

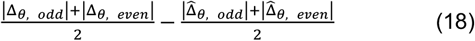

could be non-symmetrically distributed around zero. This is problematic, because a sign-flip permutation test assumes a symmetric zero mean null-distribution [31].

We avoided this problem using an intermediate transformation step. For each voxel and *θ*, we concatenated 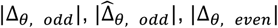, and 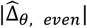 values of all subjects, and used histogram matching to transform the resulting distribution to a zero-mean normal distribution with unchanged variance. We call these new values 𝒩{|Δ_*θ, odd*_|}, 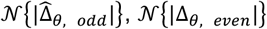, and 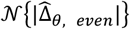. Next, similar to equation 18, for each voxel and subject, we combined these variables by averaging and taking the difference, as

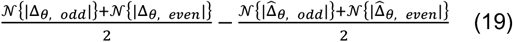

Under the null hypothesis, this new variable has a symmetric zero-mean distribution across subjects (see also *Simulation* below). Finally, we performed a sign-flip cluster permutation test on the population-level t-statistic of this variable.

In another cluster permutation test, we repeated the above analysis, comparing 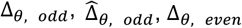, and 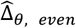, i.e. not taking the absolute values.

### Simulation

To ensure that the employed statistical procedure (equation 19) did not lead to any false positives, we performed a simulation in which we generated a different level of statistical dependency between |Δ_*θ, odd*_| and |Δ_*θ, even*_|, compared to the statistical dependency between 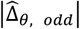 and 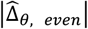. More specifically, under the null-hypothesis, we simulated an extreme case, in which we assigned independent random values to |Δ_*θ, odd*_| and |Δ_*θ, even*_| (i.e. zero dependency), but set values of 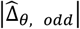 equal to values of 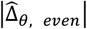 (i.e. full dependency).

To this end, for each subject and voxel, we constructed a vector of 16 elements by concatenating power values of all odd and even composite signals of all conditions (i.e. 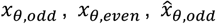, and 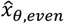, for 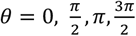). Then, for each subject, we calculated the mean of each voxel’s vector, and also their covariance matrix across the brain. Next, we drew random samples from a multivariate normal distribution with the empirically measured mean and covariance, and assigned them to 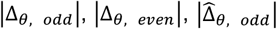. Importantly, we set 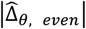 equal to 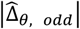 to induce the extreme case of full dependence between them. We applied the same procedure for all subjects, and, finally, applied the same cluster test to these randomly generated dataset as in the main dataset (equation 19).

We repeated this procedure 1000 times and tested if the resulting p-values were distributed uniformly. Indeed, a two-sample Kolmogorov-Smirnov test did not yield a significant difference between the distribution of p-values and a uniform distribution (P > 0.1). Thus, we concluded that the employed statistic was unbiased and did not inflate the false positive rate.

### Spectral profile of the interaction effect

To examine the spectral profile of the observed interaction between tACS and visual flicker, we computed the same test statistic that revealed the significant 10 Hz cluster (Eq. 19) at all frequencies up to 100 Hz with a spectral resolution of 1.25 Hz. For each frequency, we averaged the resulting statistic across all voxels within the 10 Hz cluster (Fig. 6a) to obtain a spectrum of interaction strength.

To assess whether any frequency other than 10 Hz showed a significant effect, we performed a sign-flip permutation test (1000 permutations). For each permutation, we randomly flipped the sign of each subject’s spectrum, averaged the spectra across subjects, and recorded the maximum value across all frequencies except 10 Hz. These 1000 maximum values formed the null distribution, controlling for multiple comparisons across frequency. For each frequency, we then compared the observed statistic against this null distribution to derive a p-value. This test assessed whether the empirical spectrum exceeded zero beyond chance level, indicating frequency-specific interaction effects.

### Heartbeat

To compare the heartbeat rate across conditions, we identified R-peaks of ECG signals and calculated inter-R-peak intervals in seconds. We assigned each interval-value to all time points between two corresponding successive R-peaks. This provided us with a continuous inter-heartbeat interval signal. Then, for each condition and each subject, we took the average across the first second (i.e., during stimulation) of all trials. For each subject, we z-scored the resulting 7 values. To normalize the data at the population level, we further divided these z-scores by their standard deviation across conditions and subjects and transformed them using a sigmoid function 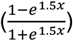. A Kolmogorov-Smirnov test did not reveal any significant difference between the distribution of the transformed data and a normal distribution (p = 0.34).

Finally, we employed a repeated measure ANOVA to compare the transformed inter-heartbeat intervals of different conditions. Specifically, we applied a 2-way repeated measure ANOVA with factors tACS phase (0 or *π*), visual flicker (present or absent), and their interaction across all conditions with tACS (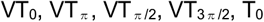, and T_π_).

### Analysis Software

All data analyses were performed in Matlab (MathWorks) using custom scripts and the open source toolbox Fieldtrip [32].

### Data and materials availability

The data that support the findings of this study are available from the corresponding authors.

## Results

### Testing linear superposition versus interaction

To test whether tCS simply superimposes on intrinsic brain activity or interacts with it, it is essential to control intrinsic activity. A practical way to achieve this is to use a sensory stimulation that elicits a reliable and well-controlled brain rhythm, providing a reference state against which the effects of tCS can be compared.

At first sight, one might consider a straightforward test: add the signals recorded during sensory stimulation alone and during tCS alone, and compare this sum to the signal recorded during simultaneous sensory stimulation and tCS (i.e., as labeled in Fig. 1, compare the sum of the signals in conditions S and T with the signal in condition ST). If the two matched, this would suggest no interaction, whereas any difference would suggest a potential interaction between tCS and sensory stimulation.

**Figure 1.**
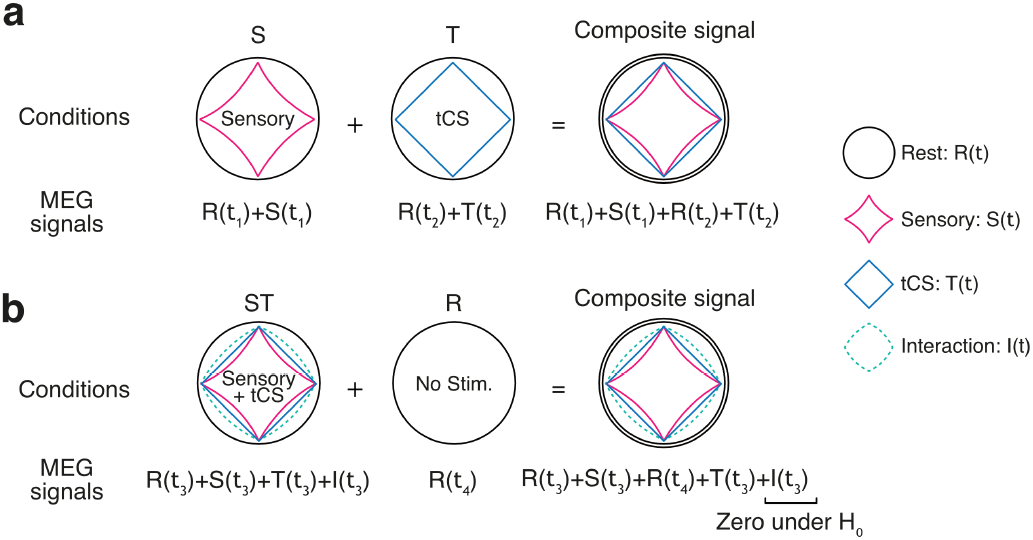
Rational of experimental approach. The illustration shows hypothetical experimental conditions and their recorded signals. Each recorded signal is modeled as the summation of a random process that models rest recordings *R*(*t*) and deviations *S*(*t*) and *T*(*t*) from the rest process that are caused by sensory stimulation and tCS, respectively. *I*(*t*) represents deviations that are caused by an interaction between sensory stimulation and tCS. Distinct subscripts for the time index (*t*) indicate that signals from different conditions are recorded at distinct times. (**a**) A composite signal constructed by summation of signals from separate sensory and tCS stimulation conditions contains two rest signals *R*(*t*) and the separate effects of tCS *T*(*t*) and sensory stimulation *S*(*t*). **(b)** A composite signal constructed by summation of signals from the simultaneous sensory and tCS stimulation condition and the rest condition contains two rest signals *R*(*t*), separate effects of tCS *T*(*t*) and sensory stimulation *S*(*t*), and potential signal *I*(*t*) that reflects an interaction between sensory stimulation and tCS. Under the null hypothesis of no interaction, the interaction term *I*(*t*) would be zero, i.e. under the null hypothesis and assuming balanced levels of tCS artifact, the composite signals in (a) and (b) would be indistinguishable.

However, this logic is flawed. The sum of sensory stimulation alone and tCS alone contains not only the separate effects of sensory and tCS stimulation, but also two copies of ongoing brain activity that is independent of stimulation (represented by the double circle for the composite signal in Fig. 1a, with one copy contributed by each condition).

By contrast, the simultaneous sensory stimulation and tCS condition contains only one copy of ongoing brain activity. In the following we will refer to this stimulation-independent ongoing brain activity as resting activity.

To make the comparison valid, we must equalize the amount of resting activity in both cases. This is achieved by adding signals from a resting condition to the simultaneous sensory stimulation and tCS signal (Fig. 1b). The result is two composite signals that are matched in resting activity, in the independent effects of sensory stimulation and tCS, and potentially in their artifact content: one representing the linear superposition of sensory stimulation and tCS applied separately (Fig. 1a), and one representing sensory stimulation and tCS applied simultaneously (Fig. 1b).

Assuming similar artifact levels, the absence of interaction yields indistinguishable composite signals, whereas any difference between them indicates an interaction between tCS and intrinsic brain activity driven by sensory stimulation. Importantly, tCS artifacts are known to interact with physiological signals such as heartbeats [12–14]. Thus, physiological signals should be compared between conditions to test if artifact levels are indeed similar between conditions.

In the following, we applied this approach in a proof-of-principle MEG experiment with 10 Hz visual flicker and 10 Hz tACS at different phase relations.

### Experiment and preprocessing

Sixteen subjects participated in the experiment, in which we recorded MEG signals during seven different stimulation conditions (Figure 2a). The experiment consisted of two elements: 1 s 10 Hz visual flicker and a 1.2 s 10 Hz tACS targeting the occipital cortex (Figure 2b). We used the 10 Hz flicker to induce steady-state brain responses with replicable phases across trials. In the context of our proposed approach, this provides defined states of intrinsic brain activity. To investigate the phase-specific interaction of tACS with intrinsic brain activity, we employed simultaneous visual flicker and tACS at four different phase differences: 0^0^, 90^0^, 180^0^, or 270^0^ (conditions: 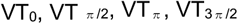, respectively). Furthermore, there were three control conditions, during which we applied either the 10 Hz visual flicker stimulus alone (V) or 10 Hz tACS alone with an initial phase of either 0° (T_0_) or 180° (T_π_). For each subject, the tACS stimulation intensity was adjusted below phosphene threshold, i.e. tACS did not induce any visual percept on its own. We ran the experiment in 1.2 s trials and randomly assigned each trial to one of the seven stimulation conditions.

**Figure 2.**
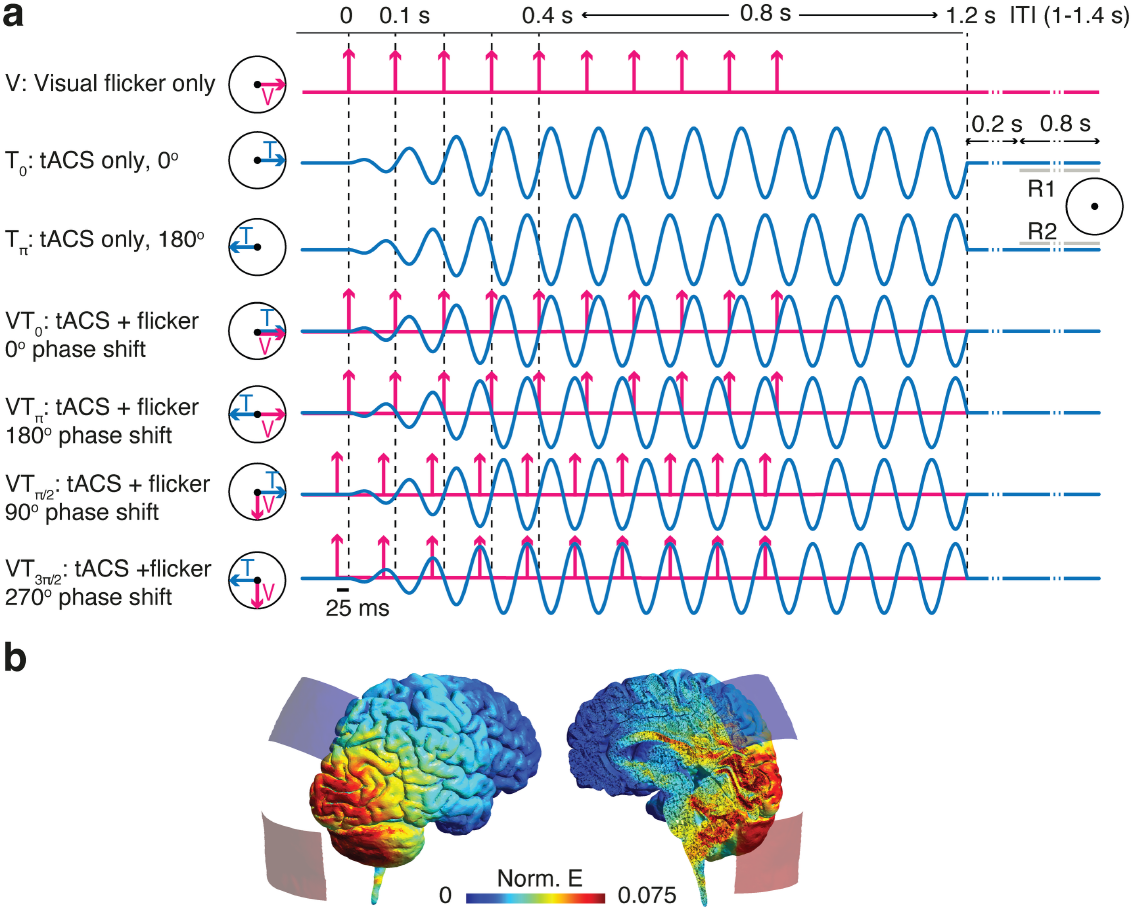
Experimental design. (**a**) The experiment consisted of 7 conditions. Each condition contained either a visual flicker stimulus at 10 Hz (V), tACS at 10 Hz (T_0_ and T_π_), or a combination of visual flicker and tACS (VT_0_, VT_π_, VT_π /2_, and 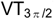). Stimulation was employed in 1.2 s trials. We discarded the first 200 ms after tACS offset to avoid offset artifacts. From each trial, we used two 800 ms segments, one during stimulation from 0.4 to 1.2 s, and one starting 0.2 s after tACS offset. We used the after-stimulation segments of T_0_ and T_π_ conditions as rest (R) signals. (**b**) Norm of electric field across the brain that results from our tACS montage at 1mA (FEM modeling using SimNIBS [27] toolbox). TACS was applied via two 35 cm^2^ rubber electrodes (two squares in the figure) that were attached such that the top edge of the first electrode lay over the midline parietal location (PZ) and the top edge of the second electrode lay over the inion (IZ).

We applied beamforming (linearly constrained minimum variance, LCMV) [28] to the MEG signals to estimate neural activity at the source level. In short, beamforming is a linear transformation that estimates the source level activity, i.e. the neural activity at different brain locations from sensor level recordings.

As a first sanity check, we compared source level activity during visual flicker (V) and during rest (R) using cluster-based permutation statistics (see Methods). We computed the pairwise phase consistency (PPC) [30] to quantify the phase alignment of neural activity across trials. As expected, 10 Hz visual flicker induced a significant increase of 10 Hz PPC in visual cortex, which reflects the steady-state response of visual cortex to the flicker stimulus (Fig. 3a, left). In contrast, 10 Hz power was significantly decreased during visual stimulation as compared to rest (Fig. 3a, right). Compared to the increase in phase-locking, the power decrease was less pronounced and excluded the primary visual area. This reflects the opposing effects that the visual stimulus had on phase-locked and non phase-locked components of alpha-band activity [33]: while presenting a visual stimulus leads to a general decrease in non phase-locked alpha-band activity, known as alpha suppression [34,35], the evoked steady-state response at the flickering frequency (10 Hz) leads to an increase in the phase-locked activity at 10 Hz. In sum, visual flicker induced the expected phase-alignment of intrinsic neural activity at the stimulation frequency.

**Figure 3.**
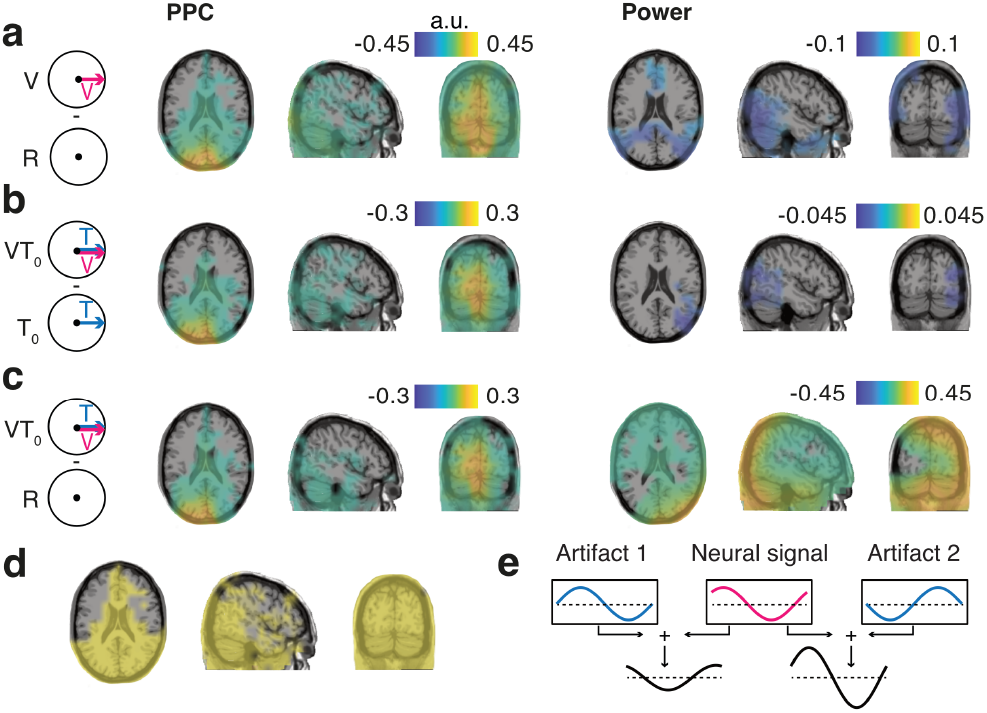
Beamforming reduces tACS artifact. (**a**) Comparing 10 Hz neural activity during visual flicker (V) and rest (R) at the source level revealed a significant increase of 10 Hz PPC in visual areas (left) and a significant decrease of 10 Hz power in occipital areas (right). Figure depicts differences in PPC and decimal log power in the source space. Pay attention that the power cluster excludes primary visual areas, because of the two opposing effects in the stimulus: while presenting a visual stimulus decreases the non phase-locked visual alpha power, a 10 Hz visual flickering increases the phase-locked 10 Hz power, as it is evident from the PPC plots. (**b**) Comparing 10 Hz neural activity during simultaneous visual flicker and tACS (VT_0_) and pure tACS (T_0_) yielded similar result as the contrast between V and R. Thus, beamforming strongly suppresses tACS artifacts. (**c**) Comparing 10 Hz neural activity of VT_0_ and rest R conditions revealed a physiologically non-plausible increase of 10 Hz power. Thus, beamformed data contained residual artifacts. (**d**) The combination of phase and power clusters resulted from comparing V and R conditions (panel a), which we considered as the region of interest (ROI) for subsequent analyses on composite signals. (**e**) Schematic illustration how residual artifacts can produce spurious power differences in the absence of true neural differences. The plot shows a 10 Hz neural signal (top middle) and two possible 10 Hz artifact signals that differ by 180 degrees in phase (top left and right). Adding these artifacts to the same neural signal results in 10 Hz signals with different amplitudes (bottom), despite identical underlying neural activity. (a.u. denotes arbitrary unit)

### Residual artifacts at the source level

Beamforming is known to suppress tACS artifacts [12]. To assess this suppression, we again compared source-level activity with and without visual flicker, but now during tACS stimulation. In other words, we compared trials with simultaneous tACS and visual flicker against trials with pure tACS at the same initial phase (VT_0_ vs. T_0_) (Fig. 3b). Because tACS intensity was below phosphene threshold, we expected similar results as for the comparison between visual flicker and rest conditions (Fig. 3a). Notably, due to the strong tACS artifacts, such a comparison does not lead to meaningful results at the sensor level (data not shown). However, at the source level we indeed observed effects in line with our prediction. There was a significant increase of occipital 10 Hz phase-locking and a significant decrease of 10 Hz power during simultaneous visual flicker and tACS conditions as compared to matched tACS-only conditions (Fig. 3b).

Previous studies have shown that, although beamforming suppresses tACS artifacts, it does not completely remove them [12,13]. We confirmed this by directly comparing the source-level activity between the combined flicker and tACS condition (VT_0_) and rest (R). Source-level activity during combined stimulation showed up to 260% higher 10 Hz power as compared to rest signals (Fig. 3c). For comparison, the 10 Hz suppression that we observed when comparing flicker only (V) and rest (R) conditions was less than 17% (Fig. 3a). Thus, the effect of tACS was more than 10-fold stronger than the effect of visual stimulation. Thus, while the power increase with tACS may in principle reflect effects on neural activity, the relative strength of this effect suggests that it mostly reflected residual tACS artifacts.

Such residual artifacts severely impede the study of tACS effects on neural activity during stimulation. For example, one might consider studying the interaction of 10 Hz tACS and visual flicker by comparing 10 Hz power between conditions with different relative phases, e.g. by comparing conditions VT_0_ and VT_π_. However, because both flicker and tACS are applied at 10 Hz in a phase-locked manner, any power difference between these two conditions could merely be due to the phase relationship of their residual artifacts. For example, as tACS in VT_0_ and VT_π_ conditions has 180 degrees phase difference (Fig. 2), their residual artifacts might also have a phase difference of 180 degrees. A linear summation of a flicker-evoked 10 Hz steady-state response and these residual artifacts leads to 10 Hz signals with different amplitudes (Fig. 3e). Therefore, even under the null-hypothesis of no interaction between neural effects of tACS and visual flicker, there would be a power difference between conditions with different relative stimulation phases. Therefore, to investigate neural effects during tACS, one needs to account for the strength and phase of residual artifacts. Next, we implement our proposed approach to isolate the desired neural effect.

### Constructing composite signals

To test for an interaction between intrinsic neural activity driven by visual flicker and tACS, we construct composite signals from simultaneous or separate stimulation conditions. Importantly, to match tACS artifacts, the two contrasted composite signals should be constructed using pairs of VT_θ_ and T_θ_ conditions that have the exact same tACS parameters (e.g. using VT_0_ and T_0_; Fig. 2a). We took additional measures to ensure that tACS artifacts did not differ between tACS conditions. First, as tACS artifacts may change during the duration of the experiment, we recorded trials of different stimulation conditions in a randomized interleaved order (Materials and Methods). Second, as tACS artifacts are known to be modulated by heart beats [12–14], we tested for heart rate differences between tACS conditions, and found no significant differences (all p-values > 0.11; repeated measure AVOVA with factors tACS phase (0 or *π*) and visual flicker (present or absent) on inter-heartbeat intervals, during tACS, and across all conditions that contain tACS). Third, to avoid any potential confound of source-reconstruction filters, we employed the same beamforming filter for all conditions. In sum, based on these measures, we assumed that, on average, MEG signals of tACS conditions with the same tACS parameters contained similar tACS artifacts.

To construct composite signals that only differ in whether they contain the potential interaction between tACS and in-phase visual flicker (VT_0_), we followed the above-described logic depicted in Fig. 1. Replacing condition S of the general case (Fig. 1) with the visual flicker V (Fig. 4), yields the following composite signals:

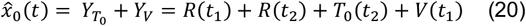

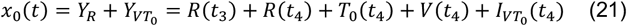

where *Y*_*C*_ is the source level activity for condition C, *V*(*t*) and *T*_0_(*t*) are the changes in the resting random process caused by the visual stimulus and 0° tACS, respectively, and 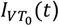 represents the interaction term. The first composite signal (20) is the summed signals from tACS alone and visual stimulation (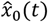, Fig. 4 left) and the other composite signal (21) is the summed signals from rest and combined stimulation (*x*_0_(*t*), Fig. 4 right). We constructed such composite signals for all four phase-differences (*x*_*θ*_(*t*) and 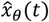, Materials and Methods).

**Figure 4.**
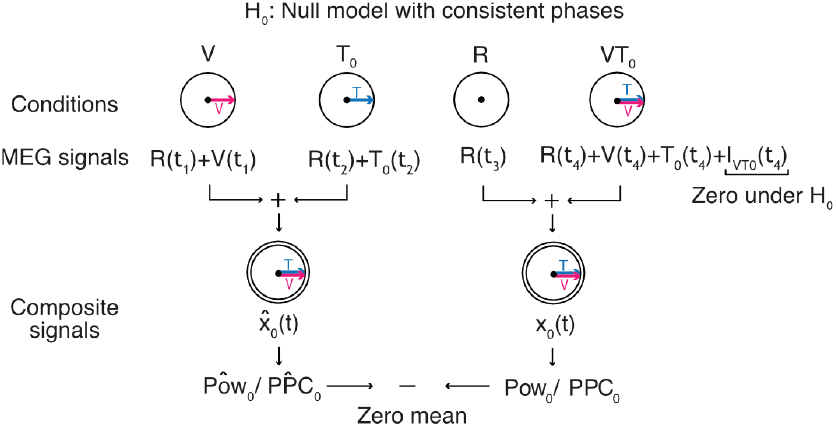
Null model with consistent phases across subjects. Under the null hypothesis, i.e. no interaction between tACS and visual stimulation, adding up neural activities of R and VT_θ_ conditions (right, *x*_*θ*_(*t*)) should lead to statistically identical signals as adding up neural activities of V and T_θ_ conditions (left, 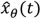, shown examples are at θ = 0). To test this, we compared power and PPC of these composite signals. However, Importantly, if such an interaction exists, this population level comparison across subjects only yields significant differences, if subjects have similar interactions at the same phaseθ; otherwise the effect may cancel across subjects in the population average.

Under the null hypothesis, the interaction term would be zero and therefore there should be no significant differences between the statistics of *x*_*θ*_(*t*) and 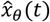. In other words, any differences between the statistics of these composite signals would indicate an interaction between tACS and intrinsic neural activity driven by visual flicker.

We applied the same cluster permutation statistic that we used for the direct contrast between conditions (Fig. 3) to compare the composite signals. Across subjects, we did not find any significant effect on power or phase-locking for either of the four relative phases between tACS and visual flicker across the brain (p > 0.096 and p > 0.2, corrected, for all power and phase comparisons, respectively). In other words, for each relative phase, the power or PPC was not affected by whether tACS was applied simultaneous to the visual stimulation (*x*_*θ*_(*t*)) or separately 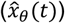.

These results could either indicate a lack of interaction between intrinsic brain activity and tACS or may merely indicate a lack of consistency across subjects. Specifically, the phase-difference, at which visual flicker and tACS interact, may vary across subjects. In this case, the effects may average out across subjects and the preformed analysis may not reveal an interaction. Thus, we next combined the composite signals into new signals that do not rely on phase consistency across subjects (Fig. 5).

**Figure 5.**
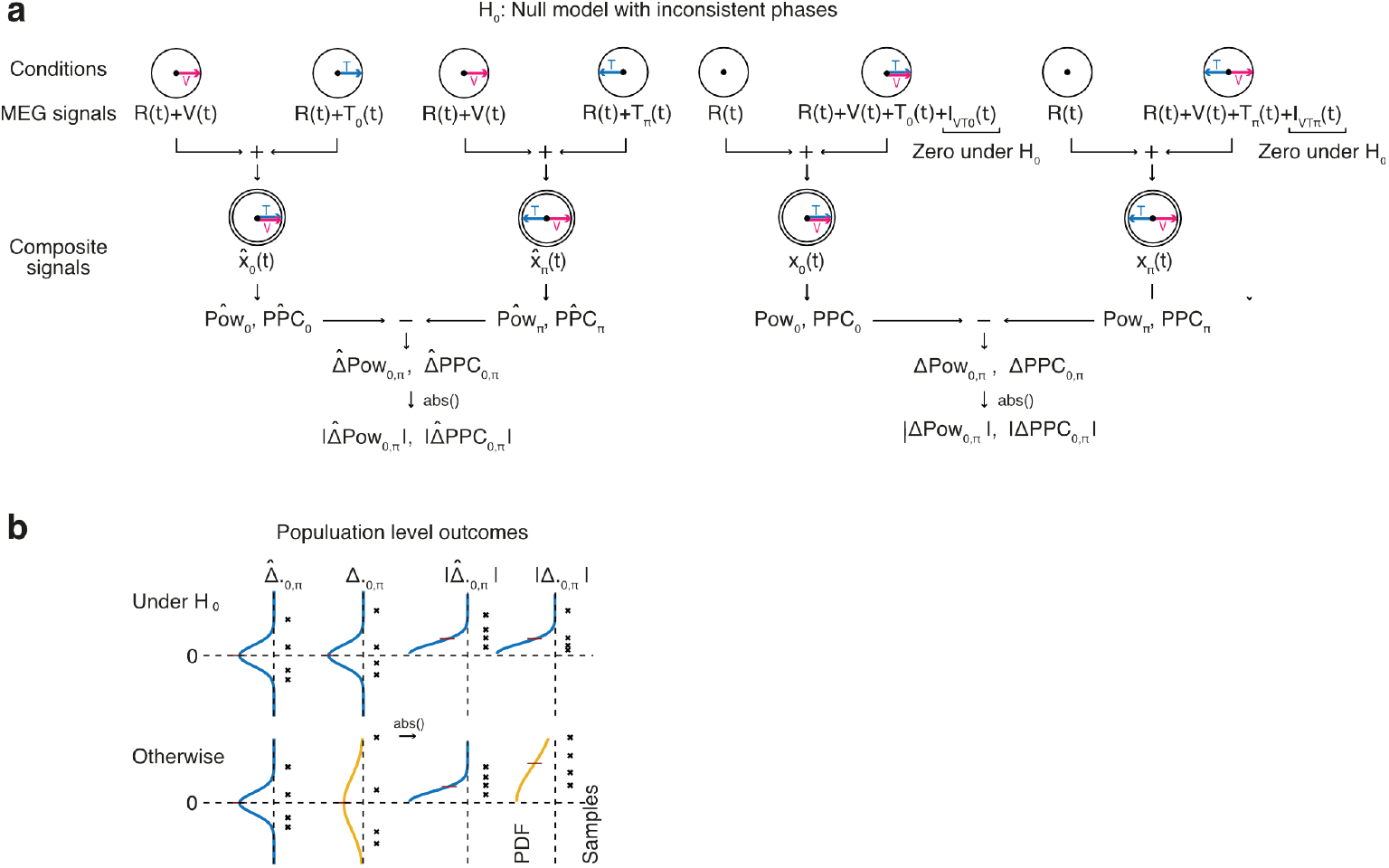
Null model with inconsistent phases across subjects. (**a**) We extended the previous model to the case, where the potential interaction between tACS and visual stimulation is not at the same θ phase across subjects. We combined anti-phase composite signals to quantify their phase dependent modulation. To this end, we calculated the absolute difference of power and PPC of ***x***_***θ***_(***t***) and ***x***_***θ***3***π***_(***t***) (|**Δ**_***θ***, ***θ***3***π***_|, right) and compared it to the absolute difference of power and PPC of 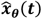 and 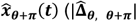, left, shown examples are at θ = 0). Note that, for simplicity and unlike in previous figures, we used the same time variable (***t***) for all conditions. (**b**) A schematic illustration of possible outcomes. Under the null hypothesis (top), i.e. no interaction between tACS and visual stimulation, both 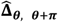 and **Δ**_***θ***, ***θ***3***π***_ (top left) and their absolute values (top right) are statistically identical. For an interaction between tACS and flicker (bottom), although mean value of

We assumed that, for each subject, independent of the phase of strongest interactions between tACS and visual flicker (*θ*), the interaction should decrease or even reverse at the opposite phase (*θ* + *π*). Therefore, we estimated how a potential interaction was modulated by phase. To this end, we computed the absolute difference of the composite signals’ power and PPC between opposite phases (Fig. 5a). If there was no interaction between neural effects of visual flicker and tACS at either *θ* or *θ* + *π*, the absolute differences obtained from *x* and 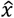 signals should be statistically equal (Fig. 5b top). Instead, if visual flicker and tACS interacted, absolute differences between opposite phases should be larger for the simultaneous stimulation (*x*) than for the separate stimulations 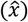 (Fig. 5b bottom).

As a first step, we applied this analysis to simulated data without any interaction (i.e. under the null hypothesis) and confirmed that the statistic was unbiased and did not produce false positives (Materials and Methods). Then, we performed this analysis on our dataset, again using the same cluster permutation statistic. This revealed an occipital cluster, in which 10 Hz power indicated an interaction between simultaneous 10 Hz tACS and visual flicker for relative phases of 0 and *π* (Fig. 6a, p = 0.032, corrected for multiple comparisons). Phase-locking (all p > 0.19, corrected) or other relative phases for power (p = 0.11, corrected) did not yield significant effects.

**Figure 6.**
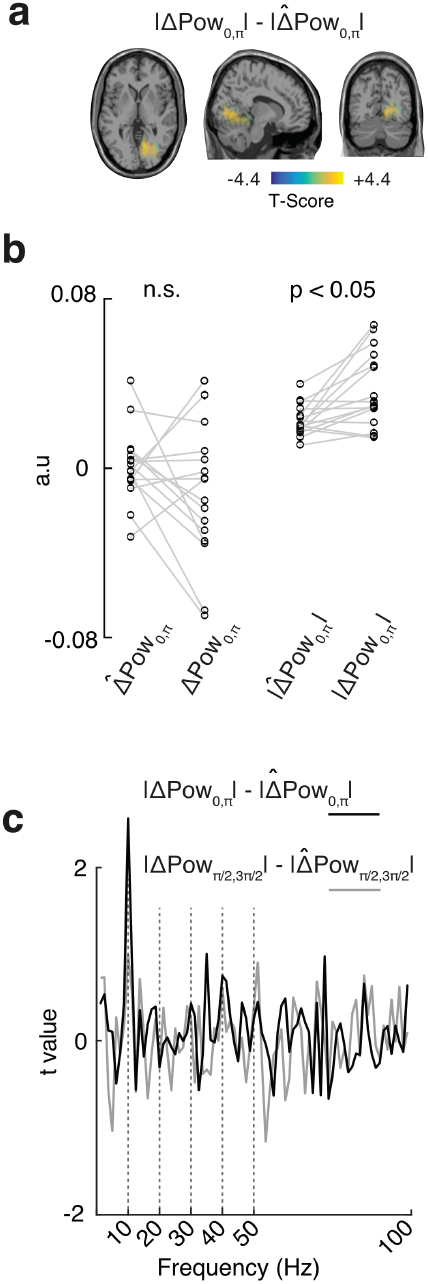
Interaction between tACS and brain activity. (**a**) Occipital cluster, where |Δ*pow*_0, *π*_| was significantly larger than 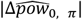. (**b**) Average activity in the identified cluster without (left) and with (right) taking the absolute. Comparing cluster averages without taking the absolute of differences (left) did not reveal any significant effect suggesting inconsistent phases of interaction across subjects. Taking the absolute revealed a significant difference (right, p = 0.032 from the original cluster-based permutation test). (**c**) Average t-value across the cluster for different frequencies. Apart from 10 Hz at 0 vs π(black line), no other frequency showed a significant average value within the cluster. (n.s.: non-significant, a.u.: arbitrary units)

To assess whether this cluster might reflect residual tACS artifacts rather than true neural modulation, we examined its spectral profile. Specifically, we tested whether the average signal within the cluster exhibited peaks at harmonic frequencies of the tACS frequency, which would suggest artifact contamination. However, no other frequency showed a significant effect within the cluster (Fig. 6c, all p > 0.3; Materials and Methods), suggesting that the observed effect was not driven by residual artifacts. In sum, two of the anti-phase conditions involving simultaneous tACS and visual flicker (i.e. VT_0_ vs VT_π_) revealed power differences that could not be explained without assuming an interaction between the neural effects of tACS and visual flicker. This provides evidence for a phase-dependent interaction between tACS and intrinsic neural activity driven by visual flicker.

We further directly investigated whether the absolute preferred phase of interaction was consistent across subjects. To this end, we removed the absolute value operator, and compared power and PPC between opposite phases for the identified occipital cluster. This did not yield any significant difference for either power or PPC (all p > 0.19, Fig. 6b, left). Moreover, in another analysis, we removed the absolute value operator and re-ran the analysis across the entire source-space. Again, we did not find any significant cluster for either power or PPC (all p > 0.14). Therefore, in agreement with the results of the first approach (Fig. 4), we concluded that the phase dependency of the interaction between tACS and visual flicker was subject specific.

In principle, tACS effects may outlast the stimulation period. Thus, in a final analysis, we tested if the observed interaction effect was detectable after stimulation offset. To this end, we performed the same cluster permutation analysis on 800 ms intervals starting 200 ms after stimulation offset. The 200 ms offset ensured that intervals were free of tACS offset artifacts. Because this interval was artifact free and to increase the sensitivity of our analysis, we simply compared VT conditions with opposite phases (i.e. VT_0_ vs VT _π_ and VT_π /2_ vs 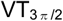). None of the tests on either power or phase yielded a spatial cluster with a significant difference (all p-values > 0.05). Therefore, we concluded that the observed interaction between tACS and intrinsic brain activity was restricted to the stimulation period.

## Discussion

Here, we present a new approach to account for stimulation artifacts and study neural effects of tCS. We propose to control intrinsic brain activity with sensory stimulation and test whether neural activity during simultaneous sensory stimulation and tCS matches the linear superposition of the effects of each applied alone. Instead of attempting to obtain artifact-free signals, we construct and compare composite signals that contain similar levels of tCS artifact but potentially differ in the interaction between tCS and intrinsic brain activity. We applied this approach in a proof-of-principle MEG experiment with 10 Hz visual flicker and 10 Hz tACS. We employed beamforming to estimate source level activity and found a phase-dependent interaction between tACS and intrinsic neural activity driven by visual flicker.

### Composite signals and residual artifacts

Our proposed approach requires that the composite signals being compared must contain similar levels of tCS artifacts. Because tCS artifacts cannot be recorded in isolation, we suggest an indirect strategy for addressing this prerequisite: monitoring physiological signals to compare artifact levels across conditions. TCS artifacts in EEG and MEG do not only depend on tCS parameters (e.g., montage, stimulation intensity, and frequency) but also on body impedance and head position (the latter is only relevant for MEG). Physiological processes such as heartbeat and respiration modulate these factors and, consequently, the characteristics of tCS artifacts [12–14]. Therefore, to ensure comparable artifact levels across conditions, we recommend continuously recording physiological signals during stimulation and neural recordings. In our proof-of-principle experiment, we monitored and excluded effects on heart rate. Future studies may also monitor additional physiological variables such as body impedance, head position and respiration rate.

Because the physiological state of participants influences tCS artifacts, it is important to avoid any temporal bias in the order of condition presentation. In our proof-of-principle study, we therefore optimized the sequence in which the seven experimental conditions were applied. Specifically, we randomized trials in successive blocks, with each block containing one trial from each condition. This protocol ensured that slow changes in tCS artifacts, such as those caused by gradual head movement or slow physiological fluctuations [12], would affect all conditions equally and minimize systematic bias due to condition order.

### Retina and the stimulation amplitude

In the presented study, we kept the tACS amplitude below individual phosphene thresholds to minimize retinal stimulation. This resulted in an average stimulation amplitude of 0.4 ± 0.16 mA (mean +/-std, 0.8 peak to peak), which translates to induced cortical electrical fields of about 0.16 V/m [36]. Although this is below the intensities often used in tACS studies [36,37], previous invasive studies have shown that even at such low stimulation intensities network level modulations are observable. For example, Reato et al. [38] observed robust entrainment effects in brain slices exposed to electric fields as low as 0.2 V/m, and Huang et al. [39] reported significant effects in the ferret brain at stimulation intensities as low as about 0.11 V/m, when applied at individual alpha frequencies. Therefore, our findings are consistent with previous studies using weak tCS intensities. Increasing the stimulation amplitude in future experiments may enhance the neural interaction effects, potentially leading to the identification of stronger and more widespread effects.

The comparatively weak stimulation and absence of phosphene perception minimized, but did not completely rule out, retinal tCS effects. In fact, the employed tACS montage is not optimally focal and may also stimulate the retina [4,40]. Future studies employing more focal stimulation montages are required to maximize the ratio of cortical to retinal stimulation [41–43,27,44] and to clarify a potential retinal contribution to the interaction between tACS and intrinsic brain activity.

### Nonlinearities in brain function

The observed interaction between the neural effects of tACS and visual flicker imply a non-additive interaction between neural effects of tACS and visual flicker. Such nonlinearities have been reported in several previous tCS studies [6,16–25]. One of the early examples of such interactions shows that the effect of transcranial direct current stimulation (tDCS) on motor performance depends on its timing relative to motor training [24]. Such dependencies have been also shown for tACS. For example, keeping the eyes closed during tACS decreases the aftereffects of occipital alpha tACS on alpha power [23]. In another study, Lustenberger et al. have shown that application of tACS locked to sleep spindles enhances motor memory consolidation [20]. Such interactions may be particularly important for tACS, for which only a small fraction of the applied currents effectively reaches the brain [41]. Therefore, it is critical to establish circumstances, under which tACS currents show their maximal effectiveness [45]. In the current study, we employed visual flicker to provide such circumstances. This approach could be readily extended to other brain areas and sensory stimulation protocols.

The nervous system shows a broad spectrum of non-linear behaviors from spike generation and synaptic integration to network interaction effects. In principle, any of such non-linearities may underlie the observed interaction between tACS and intrinsic neural activity. For example, without visual stimulation, weak tACS currents may induce sub-threshold membrane potential fluctuation with only weak effects on spiking activity. Visual flicker may bring neurons closer to their firing threshold, such that simultaneous tACS currents would induce stronger effects on spiking activity than without visual stimulation. Computational modeling and further studies – in particular invasive ones – are required to clarify the mechanisms underlying the observed interaction.

It is worth noting that we did not observe any tACS effects directly after stimulation offset. This is compatible with other previous studies that report stimulation effects restricted to the stimulation period [8,46]. For the present experiments, the lack of long-lasting effects may also be due to the relatively weak and short stimulation protocol. Indeed, a recent study that applied longer tACS and visual flicker stimuli as compared to the present experiments, found a similar interaction after tACS offset, which manifested in a lasting modulation of steady-state responses to visual flicker [47].

### Future directions

The present study establishes a basis for several avenues of future research. First, as outlined above, future work should measure a broader set of physiological parameters and use more focal stimulation montages to better rule out artifact confounds and clarify retinal contributions. It will be important to replicate our proof-of-principle findings under these conditions. Second, varying stimulation parameters could help to further elucidate the origins of the observed effects. For instance, increasing stimulation amplitude may yield more pronounced interactions, and dissociating the frequencies of tACS and sensory stimulation would provide a valuable control. For example, no interaction is expected when the tACS and visual flicker frequencies lack a harmonic relationship (e.g., tACS at 10 Hz and flicker at 25 Hz).

## Conclusion

In summary, our work xestablishes a new strategy to study online neural effects of tACS and provides direct evidence for a state-dependent and subject-specific interaction of tACS with intrinsic brain activity.

## Acknowledgements

This study was supported by the European Research Council (ERC) StG 335880 and CoG 864491 (M.S), and the Centre for Integrative Neuroscience (DFG, EXC 307) (M.S.).

## Author contributions

Conceptualization: M.S., N.N.; investigation: N.N., F.D.; formal analysis: N.N.; writing – original draft preparation: N.N.; writing – review and editing: N.N., M.S.; supervision: M.S., N.N.; resources: M.S.; funding acquisition: M.S.

